# The genetic basis of mutation rate variation in yeast

**DOI:** 10.1101/338723

**Authors:** Liangke Gou, Joshua S. Bloom, Leonid Kruglyak

## Abstract

Mutations are the root source of genetic variation and underlie the process of evolution. Although the rates at which mutations occur vary considerably between species, little is known about differences within species, or the genetic and molecular basis of these differences. Here we leveraged the power of the yeast *Saccharomyces cerevisiae* as a model system to uncover natural genetic variants that underlie variation in mutation rate. We developed a high-throughput fluctuation assay and used it to quantify mutation rates in natural yeast isolates and in 1040 segregant progeny from a cross between BY, a lab strain, and RM, a wine strain. We observed that mutation rate varies among yeast strains and is highly heritable (H^2^=0.46). We performed linkage mapping in the segregants and identified four quantitative trait loci (QTLs) underlying mutation rate variation in the cross. We fine-mapped two QTLs to the underlying causal genes, *RAD5* and *MKT1*, that contribute to mutation rate variation. These genes also underlie sensitivity to the DNA damaging agents 4NQO and MMS, suggesting a connection between spontaneous mutation rate and mutagen sensitivity.

## Introduction

Mutations are permanent changes to the genome of an organism that can result from DNA damage that is improperly repaired, from errors in DNA replication, or from the movement of mobile genetic elements. Mutations give rise to genetic variants in populations and are the wellspring of evolution. Mutations also play a major role in both inherited diseases and acquired diseases such as cancer.

The mutation rate can be defined as the number of mutational events per cell division, generation, or unit of time (Baer *et al.* 2007). Mutation rates tends to be approximately 10^−9^ to 10^−10^ mutations per base pair, per cell division, for most microbial species (Drake *et al.* 1998), making them difficult to measure and compare across individuals. As a consequence, the effects of genetic background differences on mutation rates have only been investigated on a small scale (Demerec 1937). Two types of experimental approaches have been used to measure mutation rates in yeast. The first is the fluctuation assay (Luria and Delbrück 1943). This method requires a gene with a selectable phenotype such that loss-of-function mutations in the gene enable the mutants to grow in the corresponding selective conditions. Spontaneous mutation rate is then estimated from the distribution of mutant numbers in parallel cultures. Lang and Murray applied the fluctuation assay to *S. cerevisiae* and estimated the per-base-pair mutation rate in yeast (Lang and Murray 2008a). A second method tracks mutation accumulation during experimental evolution and uses whole-genome sequencing to estimate mutation rates (Zhu *et al.* 2014). This approach also provides information on the number, locations and types of spontaneous mutations. However, this assay requires growing the mutation accumulation lines over hundreds of generations, as well as sequencing many genomes. Although the fluctuation assay is faster and cheaper, the need for many parallel cultures makes it laborious to extend it to many different strains.

Here we developed a modified version of the fluctuation assay to enable higher-throughput measurements of spontaneous mutation rates. We used the new assay to quantify mutation rates across genetically distinct yeast strains and observed considerable variation. To find the genes underlying the observed variation, we applied the modified fluctuation assay to a large panel of 1040 segregants from a cross between the laboratory strain BY4724 (hereafter referred to as BY) and the vineyard strain RM11-1a (hereafter referred to as RM). We identified four loci associated with mutation rate variation and narrowed the two loci that contributed the most to mutation rate variation to missense variants in the genes *RAD5* and *MKT1*. We also found interactions between alleles of *RAD5* and *MKT1*.

## Materials and Methods

### Yeast strains and media

Seven natural *S. cerevisiae* strains (Table S1) were used in this study. The 1040 segregants derived from BY4724 (MATa) and RM11-1a (MATa, *MKT1*-BY, hoΔ::HphMX, flo8Δ::NatMX) were generated, genotyped and described previously (Bloom *et al.* 2013). The RM::*MKT1*-BY strain was made previously by our lab. The BY::*RAD5*-RM strain and the *RAD5* variants substitution strains (Table 1) were from Demogines et al (Demogines *et al.* 2008a). For fluctuation assay, yeast was grown in synthetic complete liquid medium without arginine (SC-Arg) before plating onto selective plates. For DNA damaging agents sensitivity assays, yeast were grown in rich YPD medium (1% yeast extract, 2% peptone and 2% glucose) before plating onto YPD agar plates with DNA damaging agents. SC-Arg and YPD liquid media and agar plates were made according to Amberg et al (Amberg *et al.* 2005).

**Table 1.**
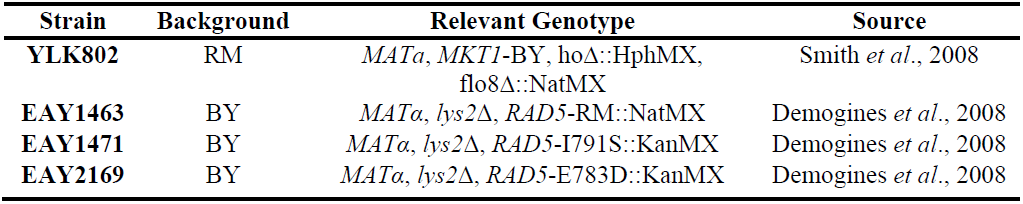
The allele replacement strains and variant substitution strains.

| Strain | Background | Relevant Genotype | Source |
| --- | --- | --- | --- |
| YLK802 | RM | *MATa*, *MKT1*-BY, *hoΔ*::HphMX,<br>flo8Δ::NatMX | Smith *et al.*, 2008 |
| EAY1463 | BY | *MATα*, *lys2Δ*, *RAD5*-RM::NatMX | Demogines *et al.*, 2008 |
| EAY1471 | BY | *MATα*, *lys2Δ*, *RAD5*-I791S::KanMX | Demogines *et al.*, 2008 |
| EAY2169 | BY | *MATα*, *lys2Δ*, *RAD5*-E783D::KanMX | Demogines *et al.*, 2008 |

### Selection agar plate construction

Selective canavanine plates were made from arginine minus synthetic complete agar medium with 60mg/liter L-canavanine (Sigma C1625). The canavanine plates were dried by incubating at 30°C overnight. Selective plates for the DNA damaging agents sensitivity assay were made with YPD agar medium containing the respective agents at the concentrations indicated in Table 2. 50ml of the agar medium was poured into each Nunc OmniTray plates (Thermo Scientific 264728) and placed on a flat surface to solidify. Each experiment was performed with the same batch of selection plates. The concentrations for 4NQO (Sigma N8141), MMS (Sigma 64382) and H_2_O_2_ (Sigma 216763) were 0.1µg/ml, 0.01% and 4mM. These concentrations capture the sensitivity difference between the segregants, while maintaining enough colony growth for QTL mapping.

**Table 2.** DNA damaging agents used for the sensitivity assay.

| **Agent** | **Agent characteristic** |
| --- | --- |
| Hydrogen peroxide (H<sub>2</sub>O<sub>2</sub>) | Altering DNA structure |
| Methyl methane sulfonate (MMS) | Altering (alkylating) DNA bases |
| 4-nitroquinoline 1-oxide (4NQO) | UV mimetic |

### Fluctuation assays

To begin the fluctuation assay, yeast were grown in synthetic complete medium without arginine (SC-Arg) in 96-well plates (Costar 3370) for ~48 hours to saturation. Saturated cultures were diluted and pinned into a new 96-well plate with liquid SC-Arg medium. This step ensured a small number of ~1000 yeast cells in the initial inoculum. Plates were sealed with a Breathe-Easy sealing membrane (Sigma Z380059) to prevent evaporation, and incubated at 30°C with shaking for ~48 hours. 100µl saturated cultures were spot-plated onto canavanine plates in a four by six configuration using a Biomek FX^P^ automated workstation. Plates with spot-plated yeast culture were dried in the laminar flow hood (Nuair) for half an hour or until dry, and incubated at 30°C for ∼48 hours. We imaged the plates using an imaging robot (S&P Robotics BM3-SC), and the number of colonies in each spot was manually counted from the images. An example of the imaged plate is shown in Figure S1.

Mutation rate was estimated using the Ma-Sandri-Sarkar Maximum Likelihood Method where the numbers of observed colonies on canavanine plates was fitted into the Luria-Delbrück distribution and a single parameter m was calculated (Sarkar *et al.* 1992). The parameter m represents the expected number of mutation events per culture. For the natural isolates and engineered strains, the mutation rate was calculated from the equation *µ* = *m*/*N*, where *N* is the average number of cells per culture (as a proxy for the number of cell divisions given the starting inoculum is much smaller than *N*). In the segregant panel, we defined a mutation rate score that was calculated as the residual phenotype after regressing out the effect of average number of cells per strain (N) from the estimate of m per strain across all of the segregants.

For each of the seven natural isolate strains, we performed ninety-six replicates of the fluctuation assay, which means we had ninety-six estimations of mutation rate. In each replicate three cultures were plated onto canavanine plates, and the number of resistance colonies in these three plates were fitted into the Luria-Delbrück distribution to estimate the mutation events per culture (m). One culture was diluted and plated onto YPD to determine the number of cells per culture (N) in each replicate. Given the number of replicates used for estimate m and N were limited, the mutation rate estimation for the seven natural isolate strains had large variance. For the BYxRM segregants panel, twelve independent replicate cultures were plated onto canavanine plates for every segregant. The number of canavanine resistant colonies in these twelve plates was fitted into the Luria-Delbrück distribution to calculate the number of mutations per culture (m), and one culture was diluted and plated on the YPD plates to determine the number of cells (N). Given only one culture was used to estimate the number of cells (N) for each segregants, which means the mutation rate estimation from the equation *µ* = *m*/*N* would be largely affected by that one measure of N. To minimize the noise driven by the measure of N, we removed the segregants with relatively less accurate measure of N from the later QTL mapping, only 843 segregants with confident measure of m and N were used. Furthermore, we defined a mutation rate score that regressed out the effect of N instead of the division. We used the mutation rate score of segregants for later QTL mapping. For each allele replacement strain (Table 1), ninety-six replicates of fluctuation analysis were performed, providing us ninety-six estimations of mutation rate. In each replicate, twelve cultures were plated onto canavanine plate to estimate the number of mutations per culture (m), and three cultures were pooled, diluted and plated on YPD plates to determine the number of cells per culture (N).

### Yeast growth measurement for DNA damaging agents sensitivity assay

The segregant panel were originally stored in 96-well plates (Costar 3370). During the DNA damaging agents sensitivity assay, individual segregants were inoculated in two plate configurations in 384-well plates (Thermo Scientific 264574) with YPD and grown for ∼48 hours in a 30°C incubator without shaking. Saturated cultures were mixed for 1min at 2,000 r.p.m. using a MixMate (Eppendorf) before pinning. The colony handling robot (S&P Robotics BM3-SC) was used to pin segregants onto selective agar plates with 384 long pins. The plates were incubated at 30°C for ∼48 hours and imaged by the colony handling robot (S&P Robotics BM3-SC). Custom R code (Bloom *et al.* 2013) was used to determine the size of each colony and the size was used as a proxy for growth in the presence of the DNA damaging agents.

### QTL mapping

In order to control for intrinsic growth rate differences and plate position effects, we normalized the traits for growth by fitting a regression for growth of the yeast that were in the same layout configuration on control plate (YPD agar plates for mutagen sensitivity assay). Residuals were used for QTL mapping. We tested for linkage by calculating logarithm likelihood ratio (LOD scores) for each genotypic marker and trait as – *n*(ln (1 – *r*^2^)/(2 ln 10)), where r is the Pearson correlation coefficient between the segregant genotypes and the segregant mutation rate or DNA damaging agents sensitivity. The threshold declaring the significant QTL effect was calculated from the empirical null distribution of the maximum LOD score determined from 1,000 permutations (Churchill and Doerge 1994). The estimated 5% family-wise error rate significance thresholds were 3.52, 3.62, 3.61 and 3.64 for mutation rate, mutagen sensitivity for 4NQO, MMS and H_2_O_2_ respectively. The 95% confidence intervals were determined using a 1.5 LOD score drop. The code and the data for QTL mapping is available at https://github.com/gouliangke/Mutation-rate/tree/master/qtl_mapping

### Amplicon sequencing of the *CAN1* region in segregants

1040 segregants were assigned into four groups according to their alleles at gene *RAD5* and *MKT1* (Table S2). We collected the cananvanine resistant colonies from the canavanine plates that we used to measure the mutation rate of segregants in the previous fluctuation analysis. A single canavanine resistant colony (if any) was picked from each segregant and the picked colonies from the same group were pooled together for DNA extraction. DNA was extracted from the pool using the Qiagen DNeasy Blood & Tissue Kit. The *CAN1* region was amplified from the DNA of four groups using the Phusion High-Fidelity DNA Polymerase (Thermo Fisher Scientific) and eight pairs of designed primers (File S3). The amplicon sequencing library was prepared using the Illumina Nextera DNA Library Prep Kit with the adjusted protocols to skip the nextera treatment. The library was then sequenced on the MiSeq platform using the MiSeq Reagent V2 Nano Kit. The sequencing read counts for each group were controlled to be the same (Figure S7). Custom R codes were used to detect the mutation rate spectrum of each group. The code for the mutation rate spectrum analysis is available at https://github.com/gouliangke/Mutation-rate/tree/master/mutation_spectrum

## Results

### High-throughput fluctuation assay for measuring mutation rates

The fluctuation assay for measuring mutation rate involves growing many parallel cultures, each starting from a small number of cells, under non-selective conditions, followed by plating to selective medium to identify mutants. The number of mutations that occurs in each culture should follow the Poisson distribution, as mutations arise spontaneously. However, the number of mutant cells that survive on the selective plates can vary greatly, because early mutations are inherited by all offspring of the mutant. This leads to the “jackpot” effect, in which some cultures contain a large number of mutant individuals. The number of observed mutant cells per culture follows the Luria-Delbrück distribution (Luria and Delbrück 1943), and the Ma-Sandri-Sarkar maximum likelihood method can be used to estimate the expected number of mutations per culture from the observed numbers of mutants (Sarkar *et al.* 1992). The underlying mutation rate is then calculated by dividing the number of mutations per culture by the average number of cells per culture [1,4]. Here we measured rare spontaneous loss-of-function mutations in the gene *CAN1*, which encodes an arginine permease. Yeast cells carrying loss-of-function mutations in *CAN1* can grow on canavanine, an otherwise toxic arginine analog. Typically, fluctuation assays are labor-intensive and have limited throughput, because a large number of parallel cultures is required for estimating the mutation rate in each assay, and several replicate assays are needed for a robust measurement of the mutation rate in each strain (Lang and Murray 2008b). We modified the fluctuation assay into a high-throughput method for measuring mutation rates in many strains in parallel. We grew cultures in 96-well plates, automated the spotting of cultures, and used high-resolution imaging to rapidly count mutants on many plates (Methods, Figure 1A). The automated spotting process for 96 strains took only approximately twenty minutes, and the imaging process required even less time. These improvements enabled us to measure the spontaneous mutation rates in the hundreds of strains necessary for genetic mapping.

**Figure 1.**
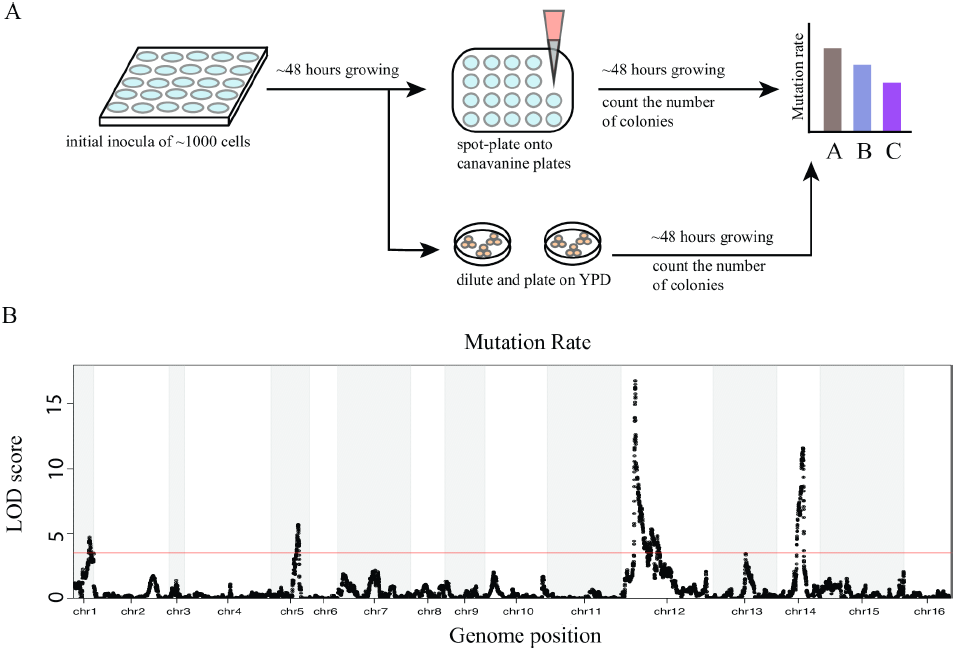
Linkage analysis identified four loci underlying mutation rate variation. (A) The fluctuation assay was performed as shown in the workflow. The assay started with a small number of cells growing in 96-well plates in liquid SC-Arg medium for ∼48 hours, followed by plating onto selective agar plates with canavanine. A proportion of the cultures were diluted to measure the number of cells per culture (Methods). Plates were imaged two days after spot-plating, and the number of colonies on canavanine plate was counted. (B) LOD score for mutation rate variation is plotted against the genetic map. The 4 significant QTLs explain 20.7% of the phenotypic variance. The red line indicates a 5% FWER significance threshold (LOD =3.52).

### Spontaneous mutation rate varies among yeast isolates

To investigate mutation rate variation among *S. cerevisiae* strains, we measured the spontaneous mutation rate of seven yeast isolates using the high-throughput fluctuation assay (Table S1). The seven strains span a large range of yeast genetic diversity (Schacherer *et al.* 2009). We found that the mutation rates of these strains range from 1.1×10^−7^ to 5.8×10^−7^ mutations per gene per generation, with a median of 1.7×10^−7^ (Table S1, Figure S2). The median mutation rate was very similar to the previously reported mutation rate at *CAN1* (Lang and Murray 2008a). In particular, the mutation rate we observed for the BY strain (1.7×10^−7^) is very similar to the previously reported rate, which was measured in strain W303 (1.5×10^−7^) (Lang and Murray 2008a), consistent with the fact that W303 shares a large fraction of its genome with BY (Liti *et al.* 2009). An analysis of variance (ANOVA) showed that strain identity explained a significant fraction of the observed variance in mutation rates (F=69.9, df=6, *p*<2×10^−16^) (Figure S2). The fraction of total variance in mutation rates explained by the repeatability of measurements for each strain, 46%, serves as an upper bound for the estimate of the total contribution of genetic differences between strains to trait variation (broad-sense heritability or H^2^). We observed that RM, a vineyard strain, had a mutation rate higher than all other strains (Figure S2).

### Four QTLs explain the majority of observed mutation rate variation

In order to find the genetic factors underlying the difference in mutation rate between BY and RM, we performed quantitative trait locus (QTL) mapping in 1040 genotyped haploid segregants from a cross between these strains (Bloom *et al.* 2013). We measured the mutation rate of each segregant using the high-throughput fluctuation assay (Methods). We estimated the fraction of phenotypic variance explained by the additive effects of all segregating markers (narrow-sense heritability) to be 30% (Methods) (Lynch and Walsh 1998). This sets an upper bound for the expectation of the total amount of additive genetic variance that could be explained with a QTL-based model. QTL mapping in the segregant panel identified significant linkage at four distinct loci (Figure 1B). At two of the QTLs, on chromosomes XII and I, the RM allele conferred a higher mutation rate, consistent with the higher mutation rate of this strain. At the other two QTLs, on chromosomes XIV and V, the BY allele conferred a higher mutation rate (Figure S3), showing that a strain with lower trait value can nevertheless harbor trait-increasing alleles. The four detected QTLs explained 20.7% of the phenotypic variance, thus accounting for 69% of the estimated additive heritability. The loci on chromosomes XII, XIV, I and V explained 8.8%, 6.1%, 3.1% and 2.6% of the variance, respectively. We tested the four identified QTLs for pairwise interactions and found a significant interaction between the QTL on chromosome XII and the QTL on chromosome XIV that explained 1% of the phenotypic variance (F=8.41, df=1, Bonferroni-corrected *p*=0.023).

### Polymorphisms in genes *RAD5* and *MKT1* underlie the major QTLs on chromosomes XII and XIV

Ten genes fell within the confidence interval of the QTL on chromosome XII. A strong candidate was *RAD5*. Previous studies showed that natural variants in *RAD5* contribute to sensitivity to the mutagen 4-nitroquinoline 1-oxide (4NQO) (Demogines *et al.* 2008a). *RAD5* encodes a DNA repair protein involved in the error-free DNA damage tolerance (DDT) pathway (Torres-Ramos *et al.* 2002; Blastyák *et al.* 2007). The DDT pathway promotes the bypass of single-stranded DNA lesions encountered by DNA polymerases during DNA replication, thus preventing the stalling of DNA replication (Bi 2015). *RAD5* plays a crucial role in one branch of the DDT pathway called template switching (TS), in which the stalled nascent strand switches from the damaged template to the undamaged newly synthesized sister strand for extension past the lesion (Bi 2015). Two non-synonymous substitutions exist between BY and RM strains in *RAD5* (Figure 2A), at amino acid positions 783 (glutamic acid in BY and aspartic acid in RM) and 791 (isoleucine in BY and serine in RM). According to Pfam alignments (Sonnhammer *et al.* 1997), *RAD5* contains a HIRAN domain, an SNF2-related N-terminal domain, a RING-type zinc finger domain, and a helicase C-terminal domain (Figure 2A). Both non-synonymous polymorphisms mapped to the helicase domain of *RAD5* (Figure 2A), and no other sequenced strains of *S. cerevisiae* contain the aspartic acid 783 and serine 791 variants that are private to the RM strain. We used protein variation effect analyzer (PROVEAN) (Choi and Chan 2015) to predict whether the two non-synonymous substitutions have an impact on the biological function of the protein. PROVEAN showed the I791S substitution (score – 5.4) might have a strong deleterious effect, while the E783D variant (score –1.8) was not predicted to have a strong effect.

**Figure 2.**
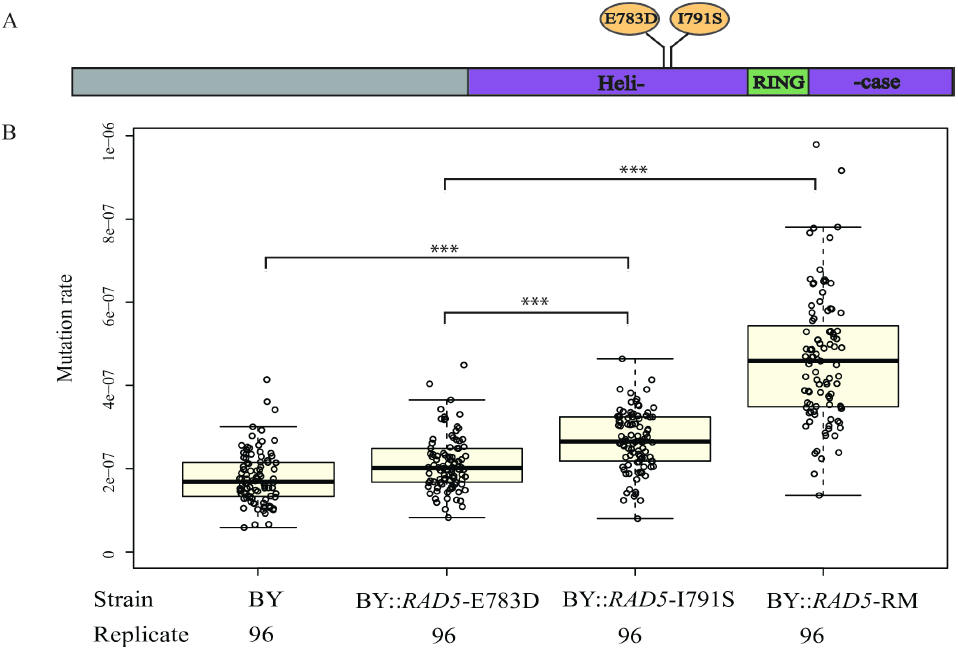
Polymorphisms in *RAD5* underlie mutation rate variation. (A) *RAD5* polymorphisms between BY and RM are located in the helicase region. The first letter for each polymorphism indicates the BY polymorphisms (E783, I791) and the second letter indicates the RM polymorphisms (D783, S791). (B) The effect of single *RAD5* polymorphism and *RAD5* whole gene replacement was tested in the BY strain background for mutation rate. For each strain, the mutation rates of ninety-six replicates were measured. Bold lines show the mean. Boxes show the interquartile range. Statistical significance was tested using a permutation t-test. Permutation p value < 0.001 is shown as ***.

Nineteen genes fell within the confidence interval of the QTL on chromosome XIV. A strong candidate was *MKT1*, which was also reported to affect 4NQO sensitivity (Demogines *et al.* 2008a). *MKT1* encodes an RNA-binding protein that affects multiple traits and underlies an eQTL hotspot in yeast (Albert and Kruglyak 2015). The RM allele of *MKT1* increases sporulation rate (Deutschbauer and Davis 2005) and improves survival at high temperature (Steinmetz *et al.* 2002), in low glucose (Parreiras *et al.* 2011), after exposure to DNA-damaging agents (Demogines *et al.* 2008a), and in high ethanol levels (Swinnen *et al.* 2012). The coding region of the BY and RM alleles of *MKT1* differs by one synonymous polymorphism and two non-synonymous substitutions. *MKT1* has an XPG domain, which is relevant to DNA repair, and an *MKT1* domain, which is related to the maintenance of K2 killer toxin (Wickner 1980). One non-synonymous variant is in the XPG domain at amino acid position 30 (aspartic acid in BY and glycine in RM), while the other non-synonymous variant is in the *MKT1* domain at position 453 (lysine in BY and arginine in RM). PROVEAN predicted a large effect of the D30G variant (score 6.7) on the function of *MKT1*, and this variant was previously found to influence sporulation rate (Deutschbauer and Davis 2005), mitochondrial genome stability (Dimitrov *et al.* 2009) and survival at high temperature [22]. The other variant (K453R) was not predicted to have a strong effect (score 0.6).

We tested whether *RAD5* and *MKT1* alleles caused differences in mutation rate by using the fluctuation test on allele replacement strains (Demogines *et al.* 2008a; Smith and Kruglyak 2008) (Table 1). The BY strain carrying the RM allele of *RAD5* (BY::*RAD5*-RM) had a higher mutation rate than the BY strain (permutation t-test, mean difference=2.9×10^−7^, p<1×10^−4^), demonstrating that the RM *RAD5* allele increases mutation rate (Figure 3A). This result is consistent with the observed difference between segregants grouped by parental allele at *RAD5* (mean difference=2.3×10^−7^). The RM strain carrying the BY allele of *MKT1* (RM::*MKT1*-BY) had a higher mutation rate than the RM strain (permutation t-test, mean difference=6.1×10^−7^, p<1×10^−4^), showing that the BY *MKT1* allele increases mutation rate (Figure 3A), consistent with the direction of effect observed in the segregants.

**Figure 3.**
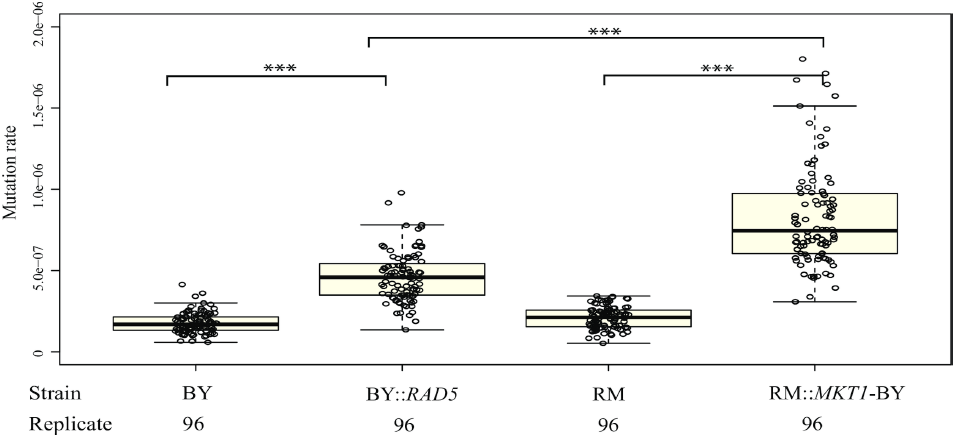
The RM allele of *RAD5* and BY allele of *MKT1* increase mutation rate. The mutation rate of two allele replacement strains, the BY strain and the RM strain are measured and compared. For each strain, ninety-six replicate measurement for mutation rate was performed. Bold lines show the mean. Boxes show the interquartile range. Statistical significance was tested using permutation t-test. Permutation p value < 0.001 is shown as ***.

To gain a finer-level understanding of the two missense variants between BY and RM in the gene *RAD5,* we tested strains (Demogines *et al.* 2008a) in which these sites in BY were individually replaced with the RM alleles (Table 1) by site-directed mutagenesis. Strains with either variant had a higher mutation rate than BY (permutation t-test, mean difference=0.9×10^−7^, p<1×10^−4^ for BY::*RAD5*-I791S; mean difference=0.3×10^−7^, p=6×10^−4^ for BY::*RAD5*-E783D) (Figure 2B), suggesting that both variants contribute to the higher mutation rate. The BY strain with the I791S substitution had a higher mutation rate than the BY strain with the E783D substitution (permutation t-test, mean difference=0.6×10^−7^, p<1×10^−4^) (Figure 2B), consistent with the PROVEAN prediction of a stronger effect for the I791S variant. However, neither variant alone nor the additive effect of the two variants fully recapitulated the increase in mutation rate that we observed when replacing the entire coding region of *RAD5* in BY with the RM allele (F=67.6, df=1, p=3.3×10^−15^), suggesting an interaction between the two variants.

### Mutation rate shares two large effect QTLs with growth on DNA damaging agents 4NQO and MMS

Deficiencies in DNA repair can increase mutation rate (Supek and Lehner 2015; Sabarinathan *et al.* 2016) and increase sensitivity to DNA damaging agents such as alkylating compounds and UV light (Sun and Moses 1991; Frankfurt 1991). We hypothesized that genetic variants that cause deficiencies in DNA repair may underlie QTLs for both mutation rate variation and sensitivity to DNA damaging agents. Previously, Demogines *et al.* identified a large-effect QTL on chromosome XII for MMS and 4NQO sensitivity in a panel of 123 segregants from a cross between BY and RM (Demogines *et al.* 2008a). Additionally, they identified a QTL on chromosome XIV for 4NQO sensitivity by using backcrossing and bulk segregant analysis. These QTLs overlapped with the major QTLs that we identified for mutation rate variation, and the underlying causal genes for 4NQO sensitivity were also *RAD5* and *MKT1*.

To follow up on these results, we measured sensitivity to three different DNA damaging agents in our panel of 1040 segregants (Table 2). The compounds assayed included methyl methanesulfonate (MMS), an alkylating agent that induces DNA double strand breaks and stalls replication forks (Hampsey 1997), 4NQO, an ultraviolet light mimetic agent (Hampsey 1997) and hydrogen peroxide (H_2_O_2_), a compound that induces DNA single and double strand breaks (Hampsey 1997). We observed that segregants with higher mutation rate, and presumably less efficient DNA repair systems, were more sensitive to MMS, 4NQO and H_2_O_2_ (Figure S4), consistent with our hypothesis that deficiencies in DNA repair increase the rate of spontaneous mutations and increase sensitivity to DNA damaging agents. We identified two large-effect QTLs for 4NQO and MMS sensitivity that overlapped with the major QTLs for mutation rate (Figure 4A and B). An interaction between *RAD5* and *MKT1* was observed for 4NQO sensitivity (F=8.5, df=1, p=0.004) (Figure S5). The QTLs on chromosome 12 and 14 were still observed in the linkage mapping for H_2_O_2_, but they had small effects (Figure S6). The large effect QTLs detected for H_2_O_2_ sensitivity on other chromosomes likely reflects trait-specific effects of variants acting on sensitivity to H_2_O_2_ (Figure S6).

**Figure 4.**
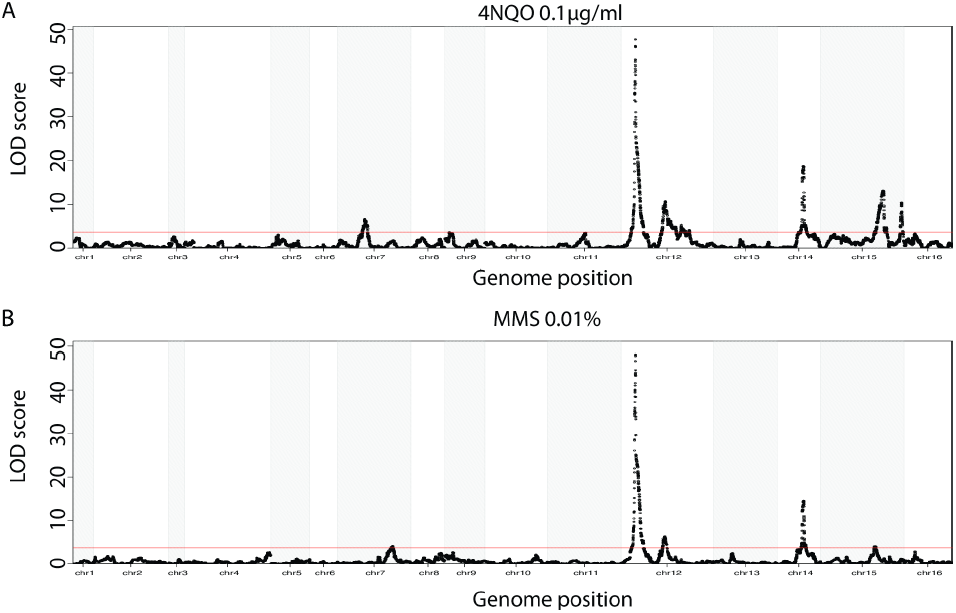
Loci underlying mutation rate variation, 4NQO sensitivity and MMS sensitivity are overlapped. (A-B) The LOD scores for 4NQO (0.1 µg/ml) sensitivity and MMS (0.01%) sensitivity are plotted against the genetic map. The red line indicates a 5% FWER significance threshold (LOD=3.62 for 4NQO and LOD=3.61 for MMS).

### Similar mutation spectra in segregants with different *RAD5* and *MKT1* genotypes

In order to gain a better understanding of how genetic variation in *RAD5* and *MKT1* might influence the DNA damage repair process, we characterized the spontaneous mutation spectrum at *CAN1* in the segregants. We divided 1040 segregants into four groups based on their genotypes at *RAD5* and *MKT1* and sequenced pools of *CAN1*-resistant mutants from each group (Table S2; Figure S7). The mutation spectra of the four groups are shown in Table S3 and Figure S8. C:G > T:A transitions were the most frequently observed mutations. A:T > G:C was the rarest transition, and A:T > T:A was the rarest transversion. The spectra for single base pair substitutions observed here (Figure S9) are similar to previous observations based on whole-genome sequencing of mutation accumulation strains (Zhu *et al.* 2014). While there were some differences in the relative frequencies of specific mutation types (for instance, more C:G > G:C transversions in segregants with the RM *MKT1* allele and more A:T > C:G transversions in segregants with BY *RAD5* allele), these mutation differences were not statistically significant after correction for multiple testing.

## Discussion

We developed and implemented a high-throughput fluctuation assay to directly measure mutation rates in yeast. We used this assay to map four QTLs that influence differences in the spontaneous mutation rate, and narrowed the two QTLs with the largest effects to causal genes and variants. We attempted to gain insight into how these variants might affect the mutation rate by comparing mutational spectra of segregants grouped by genotype, but the differences we observed did not reach statistical significance.

We identified *RAD5* as the gene underlying the QTL with the largest effect on mutation rate. *RAD5* encodes a DNA helicase and ubiquitin ligase involved in error-free DNA damage tolerance (DDT), a pathway that facilitates chromosome replication through DNA lesions (Hishida *et al.* 2009; Unk *et al.* 2010). Previous work showed that Rad5 is a structure-specific DNA helicase that is able to carry out replication fork regression (Blastyák *et al.* 2007), a process of remodeling the replication fork into four-way junctions when replication perturbations are encountered (Neelsen and Lopes 2015). This process was hypothesized to promote DNA damage tolerance and repair during replication (Neelsen and Lopes 2015). We showed that two non-synonymous variants between BY and RM in the helicase domain affect mutation rate. The RM allele of *RAD5* increases the sensitivity of yeast to 4NQO and MMS (Demogines *et al.* 2008b), probably due to a defect in replication fork regression. Thus the RM allele of *RAD5* causes both decreased growth in mutagenic conditions and a higher mutation rate in non-stressful normal conditions.

We furthermore showed that polymorphisms in *MKT1* contribute to mutation rate variation. *MKT1* is a highly pleiotropic gene that has been shown to affect levels of transcript and protein abundance for numerous genes (Smith and Kruglyak 2008) (Foss *et al.* 2011), as well as numerous cellular phenotypes (Steinmetz *et al.* 2002; Deutschbauer and Davis 2005; Demogines *et al.* 2008a; Parreiras *et al.* 2011; Swinnen *et al.* 2012; Wang and Kruglyak 2014; Albert and Kruglyak 2015). The BY and RM alleles of *MKT1* differ by two non-synonymous substitutions that map to amino acid positions 30 (aspartic acid in BY; glycine in RM) and 453 (lysine in BY; arginine in RM). The latter variant (K453R) is located in the MKT1 domain, which is required for activity of the Mkt1 protein in maintaining K2 killer toxin (Vermut *et al.* 1994). The former variant (D30G) localizes to the XPG-N (the N-terminus of XPG) domain. Four other yeast proteins contain this domain: Exo1, Din7, Rad27 and Rad2. All of these proteins have functions related to DNA repair and cellular response to DNA damage, including DNA double-strand break repair (Exo1) (Tkach *et al.* 2012), DNA mismatch repair (Exo1, Din7) (Koprowski *et al.* 2003; Tran *et al.* 2007), nucleotide excision repair (Rad2) (Northam *et al.* 2010), ribonucleotide excision repair (Rad27) (Sparks *et al.* 2012) and large loop repair (LLR) (Rad27) (Sommer *et al.* 2008). The internal XPG (XPG-I) domain, together with XPG-N, forms the catalytic domain of the Xeroderma Pigmentosum Complementation Group G (XPG) protein. The XPG protein has well-established catalytic and structural roles in nucleotide excision repair, a DNA repair process, and acts as a cofactor for a DNA glycosylase that removes oxidized pyrimidines from DNA (Clarkson 2003). In humans, mutations in the XPG protein commonly cause Xeroderma Pigmentosum, which often leads to skin cancer (O’Donovan *et al.* 1994). The aspartic acid at position 30 in the XPG domain of Mkt1 is only found in BY and related laboratory strains. We hypothesize that Mkt1 has a previously unknown function in DNA damage repair, mediated through its XPG domain.

We found that variants in *RAD5* and *MKT1* contribute to both mutation rate variation and mutagen sensitivity. These results suggest that spontaneously occurring mutations may have a similar mutation spectrum to those created by 4NQO and MMS, and are potentially repaired by the same mechanisms. Deficient DNA repair can lead to increased sensitivity to agents such as alkylating compounds and UV light (Sun and Moses 1991; Frankfurt 1991; O’Driscoll *et al.* 1999) and to higher mutation rates at sites that are less accessible to the DNA repair system (Sabarinathan *et al.* 2016). Because mutation rates can be difficult to measure, sensitivity to mutagens may serve as a useful proxy.

Recently, Jerison et al. reported heritable differences in adaptability in 230 yeast segregants from the same cross we studied here (Jerison *et al.* 2017). They measured adaptability as the difference in fitness between a given segregant (‘founder’) and a descendant of that founder after 500 generations of experimental evolution. Interestingly, *RAD5* fell within one of the QTLs found to influence adaptability. Together with our observation that *RAD5* influences mutation rate, this finding suggests that differences in mutation rate can affect the adaptability of organisms.

## Acknowledgments

We are grateful to members of the Kruglyak lab for insightful comments on this manuscript and suggestions for experiments and data analyses. We thank Meru Sadhu for helpful discussion. We would like to especially thank the Alani lab in Cornell University for the *RAD5* allele replacement and variants substitution strains.

## Author Contributions

Conceived and designed the experiments: LG JSB LK. Performed the experiments: LG. Analyzed the data: LG JSB. Wrote the paper: LG JSB LK.

